# Tandemly repeated NBPF 3mer HOR copies (Olduvai triplets) in Neanderthal AltaiNea.hg19 assembly and two novel tandem arrays of NBPF 3mer HOR repeats in complete T2T-CHM13 assembly of human chromosome 1

**DOI:** 10.1101/2023.05.15.540746

**Authors:** Matko Glunčić, Ines Vlahović, Marija Rosandić, Vladimir Paar

**Affiliations:** Faculty of Science, University of Zagreb,10000 Zagreb, Croatia; Algebra University College, 10000 Zagreb, Croatia; University Hospital Centre Zagreb (ret.), 10000 Zagreb, Croatia; Croatian Academy of Sciences and Arts, 10000 Zagreb, Croatia

**Keywords:** NBPF repeats, Olduvai triplet, higher order repeats, T2T-CHM13 assembly, Global Repeat Map GRM, evolution

## Abstract

It is known that the ∼1.6 kb NBPF repeats are human specific and contributing to cognitive capabilities, with increasing frequency in higher order repeat 3mer HORs (Olduvai triplets). From chimpanzee to modern human there is a discontinuous jump from 0 to ∼50 tandemly organized 3mer HORs. Here we investigate the structure of NBPF 3mer HORs in the Neanderthal genome assembly of Pääbo and collaborators, comparing it to the results obtained for human hg38 chromosome 1. Our findings reveal corresponding NBPF 3mer HOR arrays in Neanderthals with slightly different monomer structures and numbers of HOR copies compared to humans. Additionally, we compute the NBPF 3mer HOR pattern for the complete telomere-to-telomere human genome assembly (T2T-CHM13) by Miga and collaborators, identifying two novel tandem arrays of NBPF 3mer HOR repeats with 5 and 9 NBPF 3mer HOR copies. We hypothesize that these arrays correspond to novel NBPF genes (here referred to as NBPFA1 and NBPFA2). Further improving the quality of the Neanderthal genome using T2T-CHM13 as a reference would be of great interest in determining the presence of such distant novel NBPF genes in the Neanderthal genome and enhancing our understanding of human evolution.

## Introduction

### High-quality Neanderthal genome sequence

Impressive progress by Pääbo and coworkers in high-quality sequencing of Neanderthal’s genome has opened new avenues in studying relation of modern humans and our closest extinct relatives Neanderthals, in quest of searching “what makes us human” (Kelso J & Prufer K, 2014, Mafessoni F et al, 2020, Noonan JP & McCallion AS, 2010, Paabo S et al, 2004, Prufer K et al, 2017, Prufer K et al, 2014). The high-quality genomes *Denisova 5 AltaiNea*.*hg19* (Prufer K et al, 2014), Chagyrskaya 8 (Mafessoni F et al, 2020), and *Vindija 33*.*19* (Prufer K et al, 2017) were determined. Under the assumption that Neanderthals had the same mutation rate (1.45 × mutations per generation per base pair) (Fu Q et al, 2014) and generation time as for present-day humans (29 y), it was suggested that Chagyrskaya 8 lived ∼ 30 ky after Denisova 5, and ∼ 30 ky before Vindija 33.19 (Mafessoni F et al, 2020). An analysis of these high-quality genomes revealed significant changes in genes expressed in the striatum of the brain, indicating the potential evolution of unique functions in the Neanderthal brain (Mafessoni F et al, 2020).

### Human specific ∼1.6 kb tandem repeat units in NBPF genes

Neuroblastoma is a solid malignancy that primarily affect children and has been the focus of intense research (Maris JM & Matthay KK, 1999, Van Roy N et al, 2009). The NBPF gene family was originally identified by the disruption of one of its members in a neuroblastoma patient (Vandepoele K et al, 2008). The NBPF genes are located on human chromosome 1 and contain a repetitive structure of ∼1.6 kb tandem repeat units known as Olduvai domains (also called NBPF domains, NBPF repeats, or DUF1220 domains), which code for Olduvai proton domains (previously called DUF1220) (Fortna A et al, 2004, Heft IE et al, 2020, O’Bleness M et al, 2014, O’Bleness MS et al, 2012, Popesco MC et al, 2006, Sikela JM & van Roy F, 2017, Vandepoele K et al, 2005) involved in human brain evolution. The term Olduvai for these repeat structures is referred to as Sikela – van Roy terminology (Sikela JM & van Roy F, 2017). Alternatively, in accordance with Willard’s terminology used for tandem repeats in centromeric region of human genome (Warburton PE & Willard HF, 1996, Waye JS & Willard HF, 1987, Willard HF, 1985), the repeat units in NBPF sequences were referred to as NBPF monomers (Paar V et al, 2011).

Studies have found that the copy number of Olduvai domains is correlated with various aspects of brain function and pathology, including brain size, cortical neuron number, IQ scores, cognitive aptitude, autism, schizophrenia, microcephaly, macrocephaly, and neuroblastoma (Andries V et al, 2015, Astling DP et al, 2017, Davis JM et al, 2014, Dumas L & Sikela JM, 2009, Dumas LJ et al, 2012, Fiddes IT et al, 2019, Heft IE et al, 2020, Keeney JG et al, 2014, Mitchell C & Silver DL, 2018, Popesco MC et al, 2006, Quick VB et al, 2016, Vandepoele K et al, 2005, Zimmer F & Montgomery SH, 2015). The association between HLS Olduvai domain copy number and the human brain evolution with increased cognitive function was suggested by Sikela and coworkers (Popesco MC et al, 2006, Sikela JM & van Roy F, 2017).

Interestingly, the copy number of NBPF domains in nonhuman species generally decreases with increasing phylogenetic distance from humans, with humans having the highest number (∼300 copies) followed by great apes (∼38-97 copies), monkeys (∼48-75 copies), and non-primate mammals (∼1-8 copies), while these domains are mostly absent in non-mammalian species (Andries V et al, 2012, Dumas L et al, 2007, Heft IE et al, 2020, Keeney JG et al, 2014, O’Bleness MS et al, 2012, Popesco MC et al, 2006, Zimmer F & Montgomery SH, 2015).

In our previous research, we discovered that the intensity of NBPF copy number human specificity is much more pronounced for tandemly organized NBPF 3mer higher order repeats (HORs) than for individual NBPF repeats (Paar V et al, 2011). Specifically, we observed a copy number of 47 HORs in humans, whereas chimpanzees, gorillas, orangutans, and rhesus macaques showed zero copy number. Recent computations using higher quality sequencing, specifically the hg38 human reference genome and NC_036879.1 chimpanzee ensemble, have yielded similar results (Gluncic M et al, 2023). Based on these findings, we have hypothesized that the tandemly organized ∼4.8 kb NBPF 3mer HOR copy number may provide an additional evolutionary signature, in conjunction with the individual ∼1.6 kb NBPF primary repeat/Olduvai domain copy number effect, potentially leading to a coherent overall effect.

In this study, we aim to compare the copy number of NBPF tandemly repeated HORs in Neanderthals and humans, as well as in comparison to the complete T2T-CHM13 assembly (Altemose N et al, 2022, Miga KH & Alexandrov IA, 2021, Nurk S et al, 2022) and chimpanzee reference Pan troglodytes NHGRI_mPanTro3-v1.1-hic.freeze_pri (CM054434.1 Pan troglodytes isolate AG18354 chromosome 1, whole genome shotgun sequence). Our findings could shed light on the role of NBPF genes in human evolution, as well as on the genetic differences between Neanderthals and modern humans.

### The ∼ 171 bp alpha satellite monomers and alpha satellite nmer Higher Order Repeats (HORs) in centromeres of human genomes – a HOR prototype

Most pronounced tandem repeats in human genome are alpha satellites located in the centromeric regions of chromosomes. These repeats consist of ∼30-50% diverged ∼171 bp alpha satellite monomer units, which serve as the primary repeats (Manuelidis L, 1978, Willard HF & Waye JS, 1987). Frequently, these alpha satellites are organized into Higher Order Repeats (HORs) (Warburton PE & Willard HF, 1996). HORs are composed of monomers arranged in multimeric repeat HOR copies that are tandemly positioned. The level of divergence between HOR copies is very small, often less than 5%, which is an order of magnitude smaller than the divergence observed between neighboring monomers (Altemose N et al, 2022, Feliciello I et al, 2020, Mellone BG & Fachinetti D, 2021, Miga KH & Alexandrov IA, 2021, Paar V et al, 2007, Paar V et al, 2005, Sullivan LL et al, 2017, Uralsky LI et al, 2019, Vlahovic I et al, 2017, Warburton PE et al, 2008, Warburton PE & Willard HF, 1996, Willard HF, 1985, Willard HF & Waye JS, 1987). These primary repeats and tandem HOR repeats are commonly referred to as Willard’s terminology. The repetitive structure of alpha satellite repeats and the organization of HOR arrays have significant implications in the study of chromosome biology, genome instability, evolution, and human disease. They serve as a source of genetic and epigenetic variation, contributing to the dynamic nature of the genome(Miga KH & Alexandrov IA, 2021, Rudd MK et al, 2006).

### NBPF 3mer HORS / Olduvai triplets

In 2011, we applied our robust HOR-searching algorithm, GRM (Global Repeat Map algorithm), to the Build 36.3 human genome assembly for chromosome 1, which led to the discovery of tandemly organized ∼4.8-kb 3mer HOR copies in NBPF genes (Paar V et al, 2011). The GRM algorithm revealed that each 3mer HOR copy is composed of three ∼ 1.6 kb NBPF monomers, denoted m01, m02 and m03, respectively (Paar V et al, 2011).

In 2012, the same pattern was recognized through another method, which involved analyzing the similarity between the ∼1.6 kb Olduvai domains present in NBPF genes. This analysis revealed that the ∼1.6 kb Olduvai domains are predominantly organized in repeating triplets with minimal divergence between the triplets (O’Bleness M et al, 2014, O’Bleness MS et al, 2012). Previously referred to as HLS/DUF1220 triplets in Sikela-van Roy terminology, and consist of three domains designated as HLS1, HLS2, and HLS3, these triplets have recently been renamed Olduvai triplets (Sikela JM & van Roy F, 2017). Table 1 presents the correspondence between different names for the same repeat pattern obtained through different computational methods.

**Table 1.** Correspondence of Sikela-van Roy terminology and Willard terminology for NBPF repeats.

| **Sikela-van Roy terminology** | **↔** | **Willard terminology** |
| --- | --- | --- |
| DUF1220 domains/ Olduvai domains | ↔ | NBPF monomers |
| HLS-1 | ↔ | m1 |
| HLS-2 | ↔ | m2 |
| HLS-3 | ↔ | m3 |
| DUF1220 triplets / Olduvai triplets | ↔ | NBPF 3mer HOR copies |

Willard terminology was used generally for $n$ monomers in $n$ mer HOR copies with $n$ distinct monomers of types m1, m2, ... mn (*Warburton PE & Willard HF, 1996, Willard HF, 1985*). For the NBPF repeats, Willard terminology is applied here to the special case of $n = 3$ , with three types of ~1.6 kb NBPF monomers m1, m2, and m3. In the computational method for HOR identification, Willard terminology is characterized by identification of ~4.8 kb NBPF 3mer HOR copies (i.e., Olduvai triplets) in the first step of computation, and their constituent monomers m1, m2 and m3 are identified in the second step within HORs (Paar V et al, 2011). On the other hand, Sikela-van Roy terminology is characterized by identification of Olduvai domains (i.e., 3mer NBPF monomers) in the first step of computation, and their corresponding Olduvai triplets (i.e., 3mer HOR copies) identified on the basis of divergence among Olduvai domains.

**Table 2.** Number of canonical NBPF 3mer HOR copies in tandemly organized 3mer HOR copie (canonical and variant) in chromosome 1 of Neanderthal AltaiNea.hg19, human hg38 and complete human T2T-CHM13 assemblies (deduced from Fig. 2)

| Assembly | Gene NBPF |  |  |  |  |  | Total No. Of canonical HORs |
| --- | --- | --- | --- | --- | --- | --- | --- |
|  | 20 | 14 | 10 | 19 | A1 | A2 |  |
| Chimpanzee<br>NHGRI_mPanTro3-v1.1 |  |  |  |  |  |  | 0 |
| Neanderthal<br>AltaiNea.hg19 | 8 | 11 | 7 | 13 |  |  | 39 |
| Human<br>HG38 | 19 | 12 | 8 | 13 |  |  | 52 |
| Human<br>T2T-chm13 | 15 | 13 | 12 | 12 | 4 | 8 | 64 |

Additionally, it is worth noting that, besides the NBPF 3mer HOR, the same HOR-searching GRM computation also identified two other prominent HORs in human chromosome 1 (Paar V et al, 2011). In hornerin genes, novel quartic HORs were discovered, consisting of primary, secondary, tertiary, and quartic repeats with lengths of approximately ∼39 bp, ∼0.35 kb, ∼0.7 kb, and ∼1.4 kb, respectively (Paar V et al, 2011). These findings were fully confirmed by (Romero V et al, 2018). Moreover, in the centromeric region, the previously known centromeric alpha satellite 11mer HORs (Warburton PE & Willard HF, 1996) were also confirmed (Gluncic M et al, 2023, Paar V et al, 2011).

## Results and discussion

### Exclusively human-specific GRM diagrams

In the first step, the GRM diagrams for chromosome 1 in Neanderthal’s AltaiNea.hg19 assembly (Prufer K et al, 2014) and in recent complete human T2T assembly T2T-CHM13 (Altemose N et al, 2022, Miga KH, 2017, Miga KH & Alexandrov IA, 2021, Nurk S et al, 2022) are computed (Fig. 1a and b, respectively). These results are then compared to GRM diagrams for hg38 (NC_000001.11) assembly of the human genome and other nonhuman assemblies from Ref. (Gluncic M et al, 2023), namely the chimpanzee assembly NC_036879.1, gorilla assembly NC_044602.1, orangutan assembly NC_036903.1 and rhesus macaque assembly NC_041754.1. The GRM peak at ∼4.8 kb corresponds to the ∼4.8 kb tandemly organized canonical NBPF 3mer HOR copies (m1m2m3), which are based on the ∼1.6 kb NBPF monomers of types m1, m2, and m3.

**Fig. 1.**
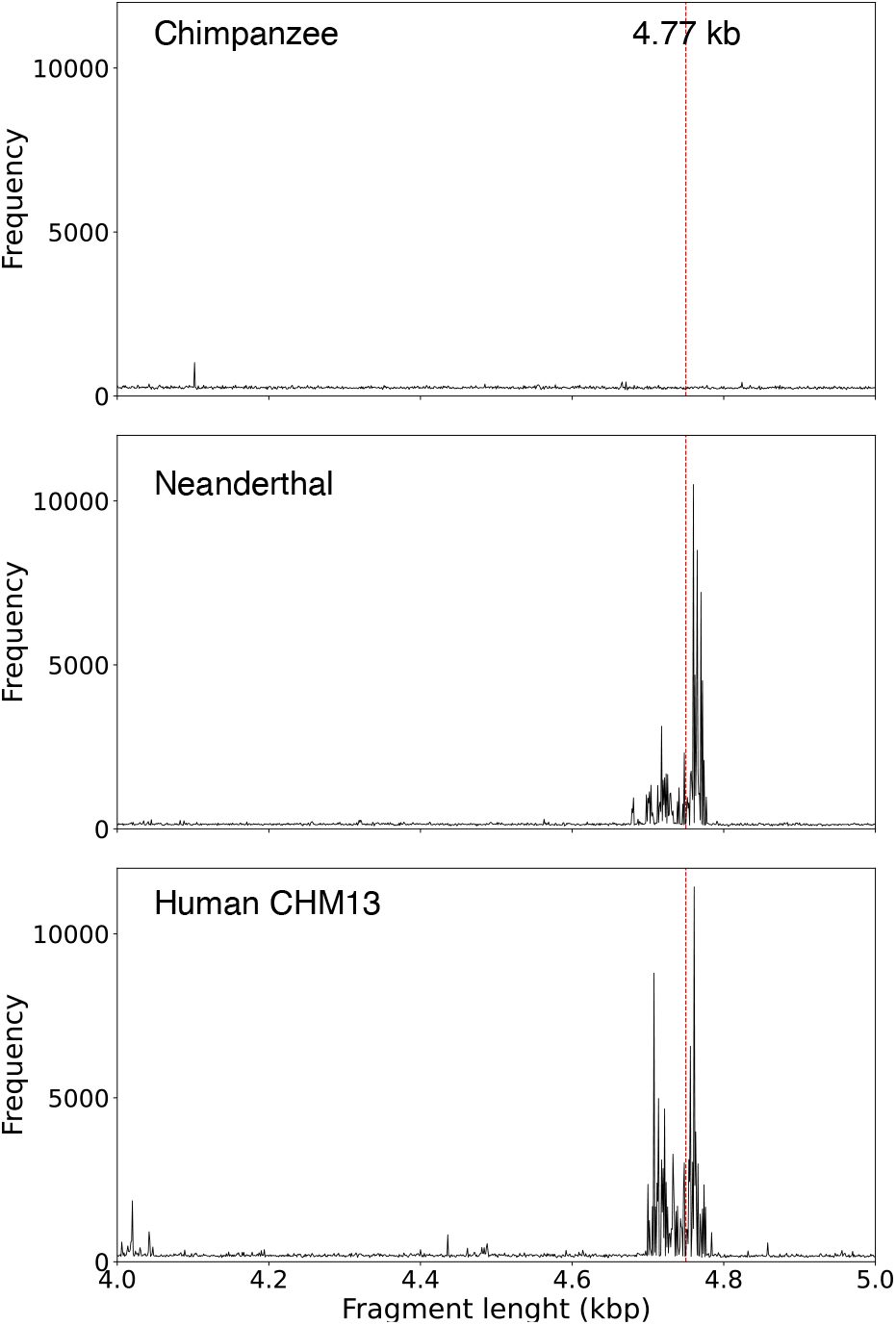
GRM diagrams for 140 Mb – 150 Mb segment of chromosome 1: a) Chimpanzee *NHGRI_mPanTro3-v1*.*1*; b) Neanderthal AltaiNea.hg19, and c) Complete human T2T CHM13 assembly. GRM diagrams have pronounced GRM peaks at ∼4.8 kb for Neanderthal and human genomes, while for chimpanzee the peak at ∼4.8 kb is absent.

### NBPF 3mer HOR copy aligned schemes

Figure 2 compares the Global Repeat Map (GRM) results for aligned NBPF 3mer higher order repeat (HOR) copies, including canonical and variant copies, in chromosome 1 across different genome assemblies. The panels show the results for the chimpanzee NHGRI_mPanTro3-v1.1 assembly (1st panel), Neanderthal AltaiNea.hg19 assembly (2nd panel), human hg38 assembly (3rd panel), and complete T2T-CHM13 assembly (4th panel). To construct these aligned NBPF HOR schemes, we first present the aligned NBPF 3mer HOR copies in the human hg38 assembly (3rd panel), which are similar to previous results computed for HLS domains(O’Bleness M et al, 2014) and GRM results (Gluncic M et al, 2023, Paar V et al, 2011). Some small differences are due to variations in the computational methods used to identify primary repeats of ∼1.6 kb and/or ∼4.8 kb secondary HOR repeats, as well as the quality of the sequenced genome. In this presentation, the NBPF monomers are horizontally grouped into HOR copies (i.e., within each 3mer HOR copy – canonical or variant), which are then aligned vertically. A blank space is inserted between any two neighboring groups of tandemly organized HOR copies, treating individual isolated monomers of types m1, m2 or m3 as single-monomer HOR copies.

**Fig. 2.**
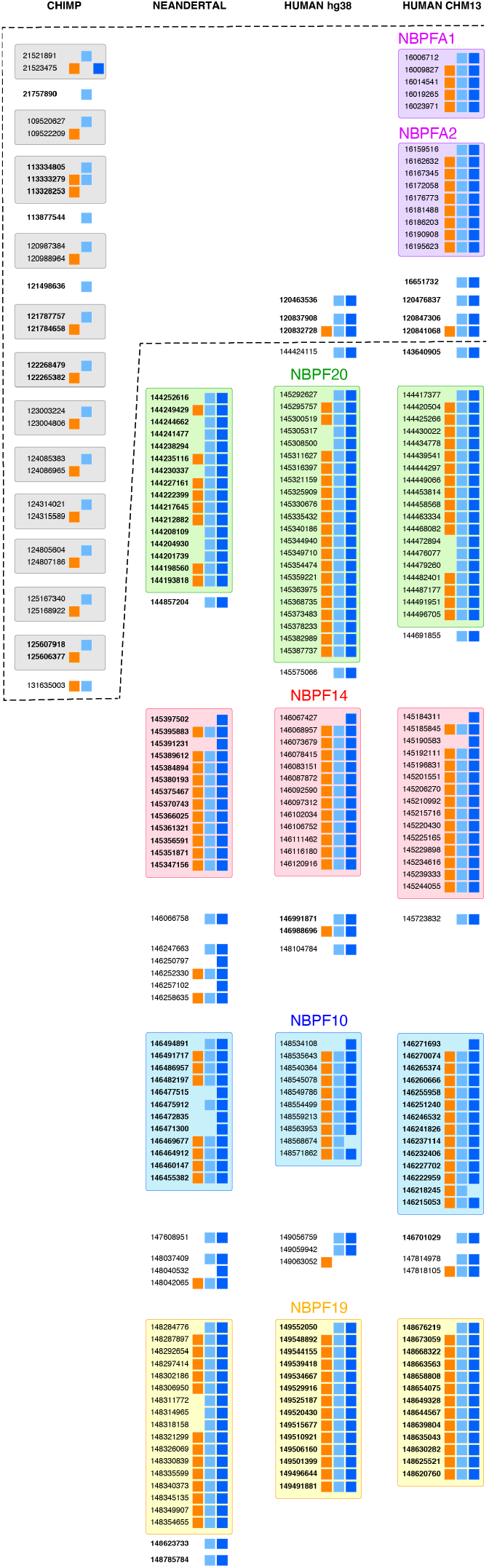
Comparison of NBPF monomer alignment scheme for NBPF 3mer HOR copies for chromosome 1 in chimpanzee *NHGRI_mPanTro3-v1*.*1* (1^st^ panel), *Neanderthal AltaiNea*.*hg19* (2^nd^ panel), *human hg38* (3^rd^ panel) and complete *human T2T-CHM13* (4^th^ panel). Boxes represent three types of NBPF monomers, denoted m1 (orange), m2 (light blue) and m3 (blue). Each row of boxes (three in canonical HOR copy, two or one in variant HOR copies) represents an NBPF HOR copy. In front of each row (HOR copy) its start position in the genomic sequence is given. Initial positions that are in bold indicate monomers that appear in the original sequence in the reverse-complement orientation. Each array of tandemly arranged HOR copies is separated from neighboring arrays and/or isolated individual monomers by blank space. Consensus monomers determined by GRM algorithm for human and Neanderthal genome are almost the same (average divergence less than 4%). A characteristics of chimpanzee NHGRI***_mPanTro3-v1*.*1*** assembly is the absence of canonical HOR copies. Moreover, except in the second variant HORcopy m1m3 (at position 21523475 bp), the m3 NBPF monomer is missing. A pronounced additional segment, identified only for the T2T-CHM13 assembly, is the appearance of two novel tandem arrays of NBPF 3mer HOR copies, the 5-copy and 9-copy arrays, interpreted as belonging to the novel genes named NBPFA1 and NBPFA2 genes.

### Human hg38 panel

The human hg38 (3rd panel) is characterized by presence of four pronounced tandemly organized arrays of NBPF 3mer HOR copies (canonical and variant). These arrays include a 22-copy array, 13-copy array, 10-copy array, and 14-copy array, respectively, with at least two neighboring canonical HOR copies arranged in tandem (Fig. 2). These arrays correspond to the four prominent NBPF genes found on human chromosome 1: NBPF20, NBPF14, NBPF10, and NBPF19, respectively. Combined, these four NBPF genes encompass a total of 59 NBPF 3mer HOR copies (52 of which are canonical). It is important to mention that the order of these four tandem arrays is reversed compared to the ordering specified in Ref. (O’Bleness M et al, 2014). To facilitate visual clarity, each tandem array is color-coded: green, red, blue, and orange, respectively. Beyond these four HOR copy arrays, there are an additional seven smaller groups of scattered HOR copies. Three of these scattered groups are positioned above the 22-copy array corresponding to the NBPF20 gene.

### Horizontal alignment of 2^nd^, 3^rd^ and 4^th^ panels

In the next step, we adjusted the relative positions of the four prominent NBPF 3mer HOR tandems in panels 2, 3, and 4 of Fig. 2. This was done by aligning the first rows of each NBPF gene horizontally while maintaining the distances within the tandem arrays. By employing this spreading technique, the first rows corresponding to the NBPF20 gene in Neanderthal AltaiNea.hg19, human hg38, and human T2T-CHM13 were horizontally aligned. Similarly, the other three notable NBPF genes (NBPF14, NBPF10, and NBPF19) were horizontally aligned as well. As a result of this spreading based on the positions of the four significant NBPF genes within each panel, the top sections of the hg38 (3rd panel) and CHM13 (4th panel) are positioned above the NBPF20 gene (green area). The top section of the 3rd panel (for hg38) exhibits three distinct sets of HOR copies (2-monomer variant HOR copy, canonical HOR copy + 2-monomer variant HOR copy, 2-monomer variant HOR copy).

### Neanderthal AltaiNea.hg19 panel

The Neanderthal AltaiNea.hg19 (2nd panel) exhibits four distinct and tandemly organized arrays of NBPF 3mer HOR copies (canonical plus variant) in a top-to-bottom manner: 16-copy, 13-copy, 12-copy, and 17-copy, respectively (Fig. 2). These arrays correspond to four prominent NBPF genes located in human chromosome 1: NBPF20, NBPF14, NBPF10, and NBPF19. Altogether, these four NBPF genes contribute to a total of 58 NBPF 3mer HOR copies (39 canonical).

When comparing the alignment of NBPF 3mer HORs between Neanderthal AltaiNea.hg19 and human hg38, the tandem HOR groups from the corresponding NBPF genes appear to have a similar order. However, it is important to consider that the Neanderthal genome was sequenced based on a comparison with an earlier human assembly, and despite its relatively high-quality sequencing, there were still gaps in the genome sequences. Notably, human hg38 has a sequencing gap of 18 Mb, which is located in proximity to the position of the NBPF20 gene. Additionally, there are several other sequencing gaps of approximately 50 kb that are further away from the NBPF genes. In the Neanderthal AltaiNea.hg19 assembly, there is a 21 Mb gap near the positions of NBPF genes, along with several other gaps of around 150 kb, 100 kb, and 50 kb close to the region of NBPF genes. Consequently, while comparing the results between Neanderthal AltaiNea.hg19 and human hg38 is reasonable, it is possible that some HOR copies have been missed in the sequencing process due to the presence of more gaps in the region near the four 3mer HOR copy-rich arrays in Neanderthal.

### Human T2T-CHM13 panel

The complete human T2T-CHM13 assembly (, represented in the 4th panel, displays four prominent and tandemly organized NBPF 3mer HOR copies that roughly align with the pattern observed in hg38 (3rd panel): a 19-copy array (green area), a 15-copy array (red), a 14-copy array (blue), and a 13-copy array (orange). This ordering of tandemly organized NBPF HOR-copy arrays roughly corresponds to the NBPF genes 20, 13, 10 and 19, respectively, similar to the hg38 case. Additionally, we identified eight scattered small groups of HOR copies outside the four major HOR copy arrays in the T2T-CHM13 sequence. Notably, above the 19-copy array (green area), there are three small, scattered groups exhibiting the same HOR pattern as observed in hg38 (2-monomer variant HOR copy, canonical HOR copy + 2-monomer variant HOR copy, 2-monomer variant HOR copy). However, at the top of the 4th panel, two additional tandemly organized arrays of NBPF 3mer HOR copies are present: a 5-mer array and a 9-mer array (violet area). We tentatively assign these NBPF tandem arrays to two genes, designated as NBPFA1 and NBPFA2, respectively. These arrays are positioned relatively close to the telomeric region, located more than 100 Mb away from the known NBPF genes. The total number of constituting NBPF 3mer HOR copies (canonical plus variant) in the T2T-CHM13 assembly is 75 (64 canonical). All NBPF genes with tandemly organized Olduvai triplets (canonical NBPF 3mer HOR copies) are located in the 1q region (O’Bleness M et al, 2014). However, several NBPF genes are also known to be located also in the 1p region, but they lack tandemly organized Olduvai triplets. For instance, in the hg38 assembly, the gene NBPF1 in 1p36.13 contains no Olduvai triplet, while the gene NBPF8 in 1p11.2 contains only one Olduvai triplet (O’Bleness M et al, 2014). The two novel NBPF genes with tandemly organized Olduvai triplets, discovered here in the CHM13 assembly and absent in hg38, are referred to as NBPFA1 and NBPFA2. As the Neanderthal AltaiNea.hg19 assembly was sequenced based on a comparison to the previous human reference genome similar to hg38, it is expected that these two novel NBPF genes are absent in the Neanderthal AltaiNea.hg19 assembly.

## Materials and Methods

The NBPF HORs were identified in the NCBI assembly (2023) of human, pan troglodytes and Neanderthal genomes using the GRM algorithm (Gluncic M & Paar V, 2013, Gluncic M et al, 2019, Vlahović I et al, 2020). The GRM algorithm is an efficient and robust method specifically designed to detect and analyze very large repeat units, such as HORs, within genomic sequences. It effectively reduces computational noise associated with detecting longer and more complex HOR repeat units, ensuring the accurate identification of peaks corresponding to HOR copies. Unlike other methods, the GRM approach directly maps symbolic DNA sequences into the frequency domain using a complete K-string ensemble, avoiding the need for statistical adjustments and local optimization of individual K-strings. This unique feature allows for straightforward identification of DNA repeats in the frequency domain without the need for mapping symbolic DNA sequences to numerical sequences. The GRM algorithm demonstrates robustness in handling deviations from ideal repeats, making it suitable for repeats with substitutions, insertions, and deletions. Additionally, it provides parameter-free identification of repeats, enabling the determination of consensus lengths and consensus sequences for primary repeats and HORs. The GRM method generates a global repeat map in a GRM diagram, identifying all prominent repeats in a given sequence without any prior knowledge of the repeats. Furthermore, once the consensus repeat unit is determined using GRM, it can be further combined with a search for dispersed HOR copies or individual constituting monomers.

## Conclussion

It is shown that the abundant tandemly organized NBPF 3mer HOR copies in complete human assembly T2T-CHM13 and in Neanderthal AltaiNea.hg19 assembly of chromosome 1 are exclusively human specific, because they are completely absent in chimpanzee chromosome 1. On the other hand, Sikela and collaborators have shown that the number of NBPF repeats, which is only about twice as high in humans compared to chimpanzees, correlates with the gradual increase in primate cognitive abilities (Davis JM et al, 2014, Dumas L & Sikela JM, 2009, Keeney JG et al, 2014). We hypothesize that the increase of cognitive abilities is coherently increased by tandemly organized NBPF higher order repeats (HORs) which are highly present in human and absent in the chimpanzee genome. It will be interesting to increase further the quality of chimpanzee and Neanderthal genome sequencing in order to fill still existing gaps in these genomes, and to test more precisely a possible coherent cognitive role of NBPF HORs.

It is interesting to note that a sophisticated phenomenon of cognitive development is related to the underlying tandem HOR pattern, which is based on the concept of DNA symmetries. This pattern represents an evolutionary trajectory characterized by symmetries, resulting in increased order and a reduction in information entropy. (Gluncic M et al, 2023, Rosandic M & Paar V, 2022, Rosandic M et al, 2022).

## Acknowledgment

This work was supported by the QuantiXLie Centre of Excellence, a project cofinanced by the Croatian Government and European Union through the European Regional Development Fund— the Competitiveness and Cohesion Operational Programme (Grant KK.01.1.1.01.0004), and the grant IP-2019-04-2757 from Croatian Science Foundation. The authors thank to Janet Kelso for discussion on Neanderthal genomics.

## Author contributions

M Glunčić: conceptualization, resources, data curation, software, formal analysis, supervision, funding acquisition, validation, investigation, visualization, methodology, project administration, and writing—original draft, review, and editing.

I Vlahović: data curation, software, and formal analysis.

M Rosandić: data curation, investigation, and visualization.

V Paar: conceptualization, investigation, visualization, methodology, and writing—original draft, review, and editing.

## References

Altemose N, Logsdon GA, Bzikadze AV, Sidhwani P, Langley SA, Caldas GV, Hoyt SJ, Uralsky L, Ryabov FD, Shew CJ, et al (2022) Complete genomic and epigenetic maps of human centromeres. Science 376: eabl. doi:10.1126/science.abl4178

Andries V, Vandepoele K, Staes K, Berx G, Bogaert P, Van Isterdael G, Ginneberge D, Parthoens E, Vandenbussche J, Gevaert K, et al (2015) Nbpf1, a tumor suppressor candidate in neuroblastoma, exerts growth inhibitory effects by inducing a g1 cell cycle arrest. BMC Cancer 15: 391. doi:10.1186/s12885-015-1408-5

Andries V, Vandepoele K, van Roy F (2012) The nbpf gene family, in: Neuroblastoma – present and future. Ed Shimada, H, InTech, Rijeka, Croatia: 185–214. doi:10.5772/28470

Astling DP, Heft IE, Jones KL, Sikela JM (2017) High resolution measurement of duf1220 domain copy number from whole genome sequence data. BMC Genomics 18: 614. doi:10.1186/s12864-017-3976-z

Davis JM, Searles VB, Anderson N, Keeney J, Dumas L, Sikela JM (2014) Duf1220 dosage is linearly associated with increasing severity of the three primary symptoms of autism. PLoS Genet 10: e100. doi:10.1371/journal.pgen.1004241

Dumas L, Kim YH, Karimpour-Fard A, Cox M, Hopkins J, Pollack JR, Sikela JM (2007) Gene copy number variation spanning 60 million years of human and primate evolution. Genome Res 17: 1266–. doi:10.1101/gr.6557307

Dumas L, Sikela JM (2009) Duf1220 domains, cognitive disease, and human brain evolution. Cold Spring Harb Symp Quant Biol 74: 375–382. doi:10.1101/sqb.2009.74.025

Dumas LJ, O’Bleness MS, Davis JM, Dickens CM, Anderson N, Keeney JG, Jackson J, Sikela M, Raznahan A, Giedd J, et al (2012) Duf1220-domain copy number implicated in human brain-size pathology and evolution. Am J Hum Genet 91: 444–454. doi:10.1016/j.ajhg.2012.07.016

Feliciello I, Pezer Z, Kordis D, Bruvo Madaric B, Ugarkovic D (2020) Evolutionary history of alpha satellite DNA repeats dispersed within human genome euchromatin. Genome Biol Evol 12: 2125–. doi:10.1093/gbe/evaa224

Fiddes IT, Pollen AA, Davis JM, Sikela JM (2019) Paired involvement of human-specific olduvai domains and notch2nl genes in human brain evolution. Hum Genet 138: 715–721. doi:10.1007/s00439-019-02018-4

Fortna A, Kim Y, MacLaren E, Marshall K, Hahn G, Meltesen L, Brenton M, Hink R, Burgers S, Hernandez-Boussard T, et al (2004) Lineage-specific gene duplication and loss in human and great ape evolution. PLoS Biol 2: E207. doi:10.1371/journal.pbio.0020207

Fu Q, Li H, Moorjani P, Jay F, Slepchenko SM, Bondarev AA, Johnson PL, Aximu-Petri A, Prufer K, de Filippo C, et al (2014) Genome sequence of a 45,000-year-old modern human from western siberia. Nature 514: 445–449. doi:10.1038/nature13810

Gluncic M, Paar V (2013) Direct mapping of symbolic DNA sequence into frequency domain in global repeat map algorithm. Nucleic Acids Res 41: e17. doi:10.1093/nar/gks721

Gluncic M, Vlahovic I, Paar V (2019) Discovery of 33mer in chromosome 21-the largest alpha satellite higher order repeat unit among all human somatic chromosomes. Sci Rep-Uk 9: doi:ARTN 1262910.1038/s41598-019-49022-2

Gluncic M, Vlahovic I, Rosandic M, Paar V (2023) Tandemly repeated nbpf hor copies (olduvai triplets): Possible impact on human brain evolution. Life Sci Alliance 6: doi:10.26508/lsa.202101306

Heft IE, Mostovoy Y, Levy-Sakin M, Ma W, Stevens AJ, Pastor S, McCaffrey J, Boffelli D, Martin DI, Xiao M, et al (2020) The driver of extreme human-specific olduvai repeat expansion remains highly active in the human genome. Genetics 214: 179–191. doi:10.1534/genetics.119.302782

Keeney JG, Dumas L, Sikela JM (2014) The case for duf1220 domain dosage as a primary contributor to anthropoid brain expansion. Front Hum Neurosci 8: 427. doi:10.3389/fnhum.2014.00427

Kelso J, Prufer K (2014) Ancient humans and the origin of modern humans. Curr Opin Genet Dev 29: 133–138. doi:10.1016/j.gde.2014.09.004

Mafessoni F, Grote S, de Filippo C, Slon V, Kolobova KA, Viola B, Markin SV, Chintalapati M, Peyregne S, Skov L, et al (2020) A high-coverage neandertal genome from chagyrskaya cave. Proc Natl Acad Sci U S A 117: 15132–1. doi:10.1073/pnas.2004944117

Manuelidis L (1978) Chromosomal localization of complex and simple repeated human dnas. Chromosoma 66: 23–32.

Maris JM, Matthay KK (1999) Molecular biology of neuroblastoma. J Clin Oncol 17: 2264–. doi:10.1200/JCO.1999.17.7.2264

Mellone BG, Fachinetti D (2021) Diverse mechanisms of centromere specification. Curr Biol 31: R1491–R. doi:10.1016/j.cub.2021.09.083

Miga KH (2017) The promises and challenges of genomic studies of human centromeres. Prog Mol Subcell Biol 56: 285–304. doi:10.1007/978-3-319-58592-5_12

Miga KH, Alexandrov IA (2021) Variation and evolution of human centromeres: A field guide and perspective. Annu Rev Genet 55: 583–602. doi:10.1146/annurev-genet-071719-020519

Mitchell C, Silver DL (2018) Enhancing our brains: Genomic mechanisms underlying cortical evolution. Semin Cell Dev Biol 76: 23–32. doi:10.1016/j.semcdb.2017.08.045

Noonan JP, McCallion AS (2010) Genomics of long-range regulatory elements. Annu Rev Genomics Hum Genet 11: 1–23. doi:10.1146/annurev-genom-082509-141651

Nurk S, Koren S, Rhie A, Rautiainen M, Bzikadze AV, Mikheenko A, Vollger MR, Altemose N, Uralsky L, Gershman A, et al (2022) The complete sequence of a human genome. Science 376: 44–53. doi:10.1126/science.abj6987

O’Bleness M, Searles VB, Dickens CM, Astling D, Albracht D, Mak AC, Lai YY, Lin C, Chu C, Graves T, et al (2014) Finished sequence and assembly of the duf1220-rich 1q21 region using a haploid human genome. BMC Genomics 15: 387. doi:10.1186/1471-2164-15-387

O’Bleness MS, Dickens CM, Dumas LJ, Kehrer-Sawatzki H, Wyckoff GJ, Sikela JM (2012) Evolutionary history and genome organization of duf1220 protein domains. G3 (Bethesda) 2: 977–986. doi:10.1534/g3.112.003061

Paabo S, Poinar H, Serre D, Jaenicke-Despres V, Hebler J, Rohland N, Kuch M, Krause J, Vigilant L, Hofreiter M (2004) Genetic analyses from ancient DNA. Annu Rev Genet 38: 645–679. doi:10.1146/annurev.genet.37.110801.143214

Paar V, Basar I, Rosandic M, Gluncic M (2007) Consensus higher order repeats and frequency of string distributions in human genome. Curr Genomics 8: 93–111.

Paar V, Gluncic M, Rosandic M, Basar I, Vlahovic I (2011) Intragene higher order repeats in neuroblastoma breakpoint family genes distinguish humans from chimpanzees. Mol Biol Evol 28: 1877–. doi:10.1093/molbev/msr009

Paar V, Pavin N, Rosandic M, Gluncic M, Basar I, Pezer R, Zinic SD (2005) Colorhor--novel graphical algorithm for fast scan of alpha satellite higher-order repeats and hor annotation for genbank sequence of human genome. Bioinformatics 21: 846–852. doi:10.1093/bioinformatics/bti072

Popesco MC, Maclaren EJ, Hopkins J, Dumas L, Cox M, Meltesen L, McGavran L, Wyckoff GJ, Sikela JM (2006) Human lineage-specific amplification, selection, and neuronal expression of duf1220 domains. Science 313: 1304–. doi:10.1126/science.1127980

Prufer K, de Filippo C, Grote S, Mafessoni F, Korlevic P, Hajdinjak M, Vernot B, Skov L, Hsieh P, Peyregne S, et al (2017) A high-coverage neandertal genome from vindija cave in croatia. Science 358: 655–658. doi:10.1126/science.aao1887

Prufer K, Racimo F, Patterson N, Jay F, Sankararaman S, Sawyer S, Heinze A, Renaud G, Sudmant PH, de Filippo C, et al (2014) The complete genome sequence of a neanderthal from the altai mountains. Nature 505: 43–49. doi:10.1038/nature12886

Quick VB, Davis JM, Olincy A, Sikela JM (2016) Duf1220 copy number is associated with schizophrenia risk and severity: Implications for understanding autism and schizophrenia as related diseases. Transl Psychiatry 6: e735. doi:10.1038/tp.2016.11

Romero V, Nakaoka H, Hosomichi K, Inoue I (2018) High order formation and evolution of hornerin in primates. Genome Biol Evol 10: 3167–. doi:10.1093/gbe/evy208

Rosandic M, Paar V (2022) Standard genetic code vs. Supersymmetry genetic code - alphabetical table vs. Physicochemical table. Biosystems 218: 10. doi:10.1016/j.biosystems.2022.104695

Rosandic M, Vlahovic I, Pilas I, Gluncic M, Paar V (2022) An explanation of exceptions from chargaff’s second parity rule/strand symmetry of DNA molecules. Genes (Basel) 13: doi:10.3390/genes13111929

Rudd MK, Wray GA, Willard HF (2006) The evolutionary dynamics of alpha-satellite. Genome Res 16: 88–96. doi:10.1101/gr.3810906

Sikela JM, van Roy F (2017) Changing the name of the nbpf/duf1220 domain to the olduvai domain. F1000Res 6:. doi:10.12688/f1000research.13586.2

Sullivan LL, Chew K, Sullivan BA (2017) Alpha satellite DNA variation and function of the human centromere. Nucleus 8: 331–339. doi:10.1080/19491034.2017.1308989

Uralsky LI, Shepelev VA, Alexandrov AA, Yurov YB, Rogaev EI, Alexandrov IA (2019) Classification and monomer-by-monomer annotation dataset of suprachromosomal family 1 alpha satellite higher-order repeats in hg38 human genome assembly. Data Brief 24: 10. doi:10.1016/j.dib.2019.103708

Van Roy N, De Preter K, Hoebeeck J, Van Maerken T, Pattyn F, Mestdagh P, Vermeulen J, Vandesompele J, Speleman F (2009) The emerging molecular pathogenesis of neuroblastoma: Implications for improved risk assessment and targeted therapy. Genome Med 1: 74. doi:10.1186/gm74

Vandepoele K, Andries V, Van Roy N, Staes K, Vandesompele J, Laureys G, De Smet E, Berx G, Speleman F, van Roy F (2008) A constitutional translocation t(1;17)(p36.2;q11.2) in a neuroblastoma patient disrupts the human nbpf1 and accn1 genes. PLoS One 3: e. doi:10.1371/journal.pone.0002207

Vandepoele K, Van Roy N, Staes K, Speleman F, van Roy F (2005) A novel gene family nbpf: Intricate structure generated by gene duplications during primate evolution. Mol Biol Evol 22: 2265–. doi:10.1093/molbev/msi222

Vlahović I, Glunčić M, Dekanić K, Mršić L, Jerković H, Martinjak I, V. P (2020) Global repeat map algorithm (grm) reveals differences in alpha satellite number of tandem and higher order repeats (hors) in human, neanderthal and chimpanzee genomes – novel tandem repeat database. 43rd International Convention on Information, Communication and Electronic Technology (MIPRO), Opatija, Croatia: 237–242. doi:10.23919/MIPRO48935.2020.9245278

Vlahovic I, Gluncic M, Rosandic M, Ugarkovic E, Paar V (2017) Regular higher order repeat structures in beetle tribolium castaneum genome. Genome Biol Evol 9: 2668–. doi:10.1093/gbe/evw174

Warburton PE, Hasson D, Guillem F, Lescale C, Jin X, Abrusan G (2008) Analysis of the largest tandemly repeated DNA families in the human genome. BMC Genomics 9: 533. doi:10.1186/1471-2164-9-533

Warburton PE, Willard HF (1996) Evolution of centromeric alpha satellite DNA: Molecular organisation within and between human primate chromosomes. In Human genome evolution, pp 121–145. BIOS Scientific Publisher

Waye JS, Willard HF (1987) Nucleotide sequence heterogeneity of alpha satellite repetitive DNA: A survey of alphoid sequences from different human chromosomes. Nucleic Acids Res 15: 7549–7569.

Willard HF (1985) Chromosome-specific organization of human alpha satellite DNA. Am J Hum Genet 37: 524–532.

Willard HF, Waye JS (1987) Chromosome-specific subsets of human alpha satellite DNA: Analysis of sequence divergence within and between chromosomal subsets and evidence for an ancestral pentameric repeat. J Mol Evol 25: 207–214.

Zimmer F, Montgomery SH (2015) Phylogenetic analysis supports a link between duf1220 domain number and primate brain expansion. Genome Biol Evol 7: 2083–. doi:10.1093/gbe/evv122

